# Epigenetic Mechanisms Inhibit Smad7 and Contribute to Fibrosis in Crohn’s Disease

**DOI:** 10.64898/2026.11.09.765430

**Authors:** Chao Li, John R. Grider, Karnam S. Murthy, Jamie Bohl, Emily Rivet, Nicole Wieghard, John F. Kuemmerle

## Abstract

**Background & Aims:** TGF-β1 is a central regulator of profibrotic gene expression, extracellular matrix deposition, and the development of intestinal fibrosis in patients with fibrostenotic Crohn’s disease (CD). Unlike the increased expression of Smad7 in mucosal and lamina propria monocytes, where it impairs Treg function, mesenchymal cells from patients with fibrostenotic CD do not exhibit elevated Smad7 expression and instead display sustained TGF-β1 signaling. We therefore sought to elucidate the mechanisms underlying the unexpected silencing of Smad7 in fibrostenotic CD.

**Methods:** Subepithelial myofibroblasts (SEMFs) and muscularis smooth muscle cells were isolated from paired normal and affected ileal tissues obtained from the same patients with each Montreal phenotype of CD, as well as from non-CD controls. Gene expression, RNA-binding affinity, transcriptional activity, promoter methylation, protein levels, and the mechanisms that regulate Smad7 expression were evaluated in ex vivo tissues and primary cultured cells. MicroRNA-21 (miR-21) localization was assessed by RNAscope. miR-21-mediated suppression of Smad7 was investigated using antagomiR-21. Smad7 silencing through promoter hypermethylation was confirmed using 5-Azacytidine treatment and DNA methyltransferase 1 (DNMT1) knockdown. Smad7 lysine acetylation and Smurf1/2-mediated ubiquitination were examined using selective inhibitors and proximity ligation assays.

**Results:** In SEMFs from patients with fibrostenotic CD, aberrant silencing of Smad7 resulted from DNMT1-dependent promoter hypermethylation and pSmad3-driven, miR-21-mediated degradation of Smad7 mRNA. In addition, Smurf1/2 promoted ubiquitination and proteasomal degradation of acetylated Smad7 protein. Inhibition of miR-21 with antagomiR-21 restored Smad7 expression and reduced TGF-β1-dependent profibrotic gene expression. Likewise, demethylation with 5-Azacytidine or knockdown of DNMT1 induced Smad7 promoter hypomethylation and restored Smad7 expression. Furthermore, modulation of the balance between p300 acetyltransferase and histone deacetylase 1 (HDAC1) stabilized Smad7 protein by protecting it from degradation.

**Conclusions:** A distinctive pattern of epigenetic silencing and post-translational regulation of Smad7 was identified in patients with fibrostenotic CD. These mechanisms collectively promote unrestrained TGF-β1 signaling by suppressing Smad7 expression and stability in SEMFs, thereby contributing to the development of intestinal fibrosis.

## INTRODUCTION

Transforming Growth Factor-β1 (TGF-β1), a multifunctional cytokine, plays two distinct roles in the intestine of patients with Crohn’s disease^1–3^. The Treg effects of TGF-β1 are critical in maintaining self-tolerance and immune homeostasis^4^. As a central mediator in the development of fibrosis, however, TGF-β1 also activates intestinal mesenchymal cells including fibroblasts, subepithelial myofibroblasts (SEMF) and smooth muscle cells (SMC) ^3,5^. Once activated, mesenchymal cells produce high levels of autocrine TGF-β1 that results in excess extracellular matrix protein production, including collagen I and fibronectin, and the increased basal proliferation rates of both that lead to the architectural distortion and scarring that characterize fibrostenosis ^1–3^. This pathogenic mechanism determines the clinical course in the 30-50% of Crohn’s patients expressing a Montreal B2 phenotype of fibrostenosis and bowel obstruction. The currently available medical therapies delay but do not prevent development of fibrostenosis and need for surgery ^6^.

Canonical TGF-β signaling is characterized by activated TGF-βRI/RII to phosphorylate receptor-based Mothers against decapentaplegic homolog 2 (Smad2) and Smad3 and their complex with common-mediator Smad4 followed by translocation to the nucleus ^4^. Smads, along with coactivators and corepressors, regulate gene transcription via the consensus Smad-binding element, CAGA box, in promoter regions of profibrotic genes including the *COL1A1* gene. Upregulation of Smad7 expression normally occurs as a negative regulator of this pathway when canonical TGF-β signaling is activated ^7–9^.

We initially observed that despite increased TGF-β1 levels and sustained Smad3 activity, Smad7 was paradoxically decreased in mesenchymal cells of affected ileum compared to normal margin in the same patient with fibrostenotic Crohn’s disease. This characteristic distinguished Montreal B2 from B1 (inflammatory) and B3 (penetrating) phenotypes ^3^. The mechanisms whereby the Smad7 response is lost despite the sustained canonical TGF-β1 signaling, however, are unknown. Mechanistically this is distinct from that which occurs in epithelial cells and lamina propria mononuclear cells where TGF-β1-dependent Treg function is inhibited because of increased Smad7 levels ^10^. The latter underlies the mechanism of action of Mongersen, an anti-Smad7 oligonucleotide in previous clinical trials for CD ^11^.

We hypothesized that epigenetic events occur in the fibrostenotic ileum that contribute to the paradoxical silencing of Smad7 in mesenchymal cells of affected ileum. Analysis of hybridization reveals an 8-mer consensus binding sequence for Homo sapiens microRNA-21(hsa-miR-21) in position 1122-1129 in the 3’ UTR of Smad7 ^12,13^. Hsa-miR-21 is implicated in the pathogenesis of IBD and other fibrotic disorders through epigenetic regulation ^14–18^. The hsa-miR-21 gene is located on the plus strand of chromosome 17q23.2 within a coding gene, *TMEM49* (vacuole membrane protein), but has its own unique promoter ^19,20^. While Smad proteins bind a conserved RNA sequence to promote pre-miRNA maturation by Drosha, the pri-miR-21 promoter region contains 3 consensus Smad3 binding sites that could regulate pri-miR-21 transcription independently of TMEM49 ^21,22^. RNA polymerase II transcribes the MIR21 gene to generate pri-miR-21, which is a hairpin-like structure. The nuclear Drosha–DGCR8 microprocessor cleaves pri-miR-21 to produce pre-miR-21, which is a 70 nt stem-loop precursor miRNA. Exportin-5 transports pre-miR-21 into the cytoplasm. Dicer processes pre-miR-21 into a mature miRNA duplex, from which miR-21-5p is typically the predominant functional strand incorporated into Argonaute ^23^.

The promoter region of Smad7 also contains several CpG islands that are targets for hypermethylation and silencing of gene expression ^24^. DNA methyltransferase (DNMT)-1, a maintenance methyltransferase, elicits and maintains hypermethylation of DNA in CpG islands ^25,26^. Hypermethylation of Smad7 by DNMT-1 has been implicated in the activation of stellate cells in the setting of liver cirrhosis ^27^. Distinctive DNA methylation patterns have been reported in IBD ^28^.

Non-epigenetic mechanisms of regulation of Smad7 protein levels that have been described were also explored. Interaction of Smad7 with the E3 ubiquitin ligases, Smad ubiquitination regulatory factors 1/2 (Smurf1/2), in the nucleus allows translocation to plasma membrane as a complex ^29,30^. This interaction permits ubiquitination and proteasomal degradation of Smad7 and activated TGF-βRI receptor ^7,31^. Post-translational acetylation and deacetylation of Smad7(K64) and Smad7(K70) govern its susceptibility to ubiquitination with acetylated Smad7 being protected from Smurf1/2-mediated ubiquitination ^32–34^.

Although the role of Smad7 has been investigated in intestinal epithelial cells and lamina propria mononuclear cells (LPMCs), its contribution to mesenchymal cells and intestinal fibrosis remains unknown. In the current study, we show that Smad7 is silenced in the affected ileum of patients with fibrostenotic CD through specific epigenetic mechanisms and post-translational modifications.

## Materials and Methods

### Isolation of subepithelial myofibroblasts (SEMFs) from patients with Crohn’s disease

Intestine was obtained from patients undergoing ileal or ileocolonic resection for CD (Supplemental material: Table 1). All specimens were obtained from patients whose Montreal phenotype had been confirmed by radiographic and pathological evaluations. . Although Montreal classification is hierarchical, comparisons in this study were made among patients with isolated fibrostenotic (B2), inflammatory (B1), or penetrating (B3) disease. Patients with concurrent fibrostenotic and penetrating disease were excluded from the analysis. Normal ileal tissue from non-CD subjects served as controls.

**Table 1.**
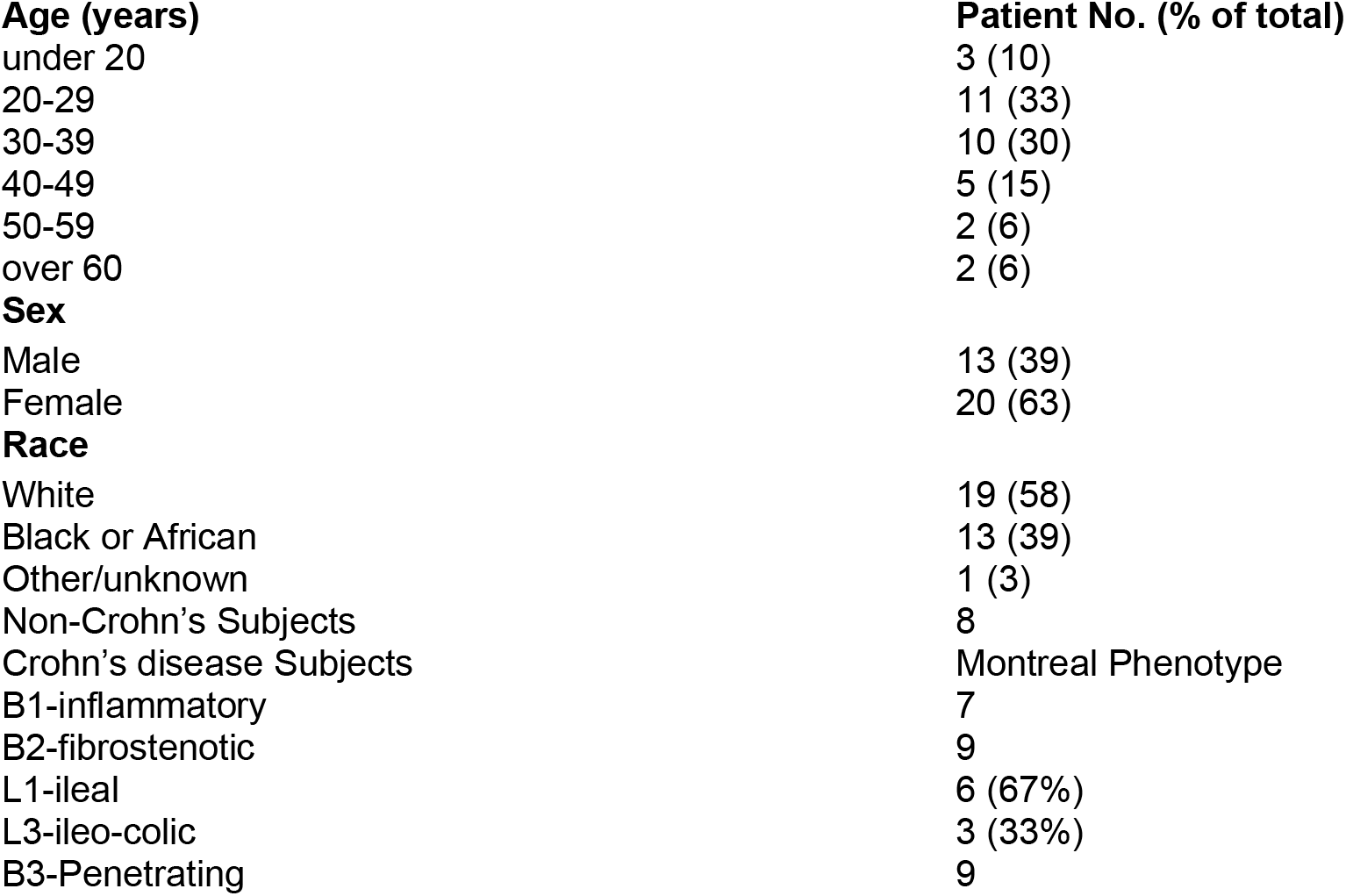
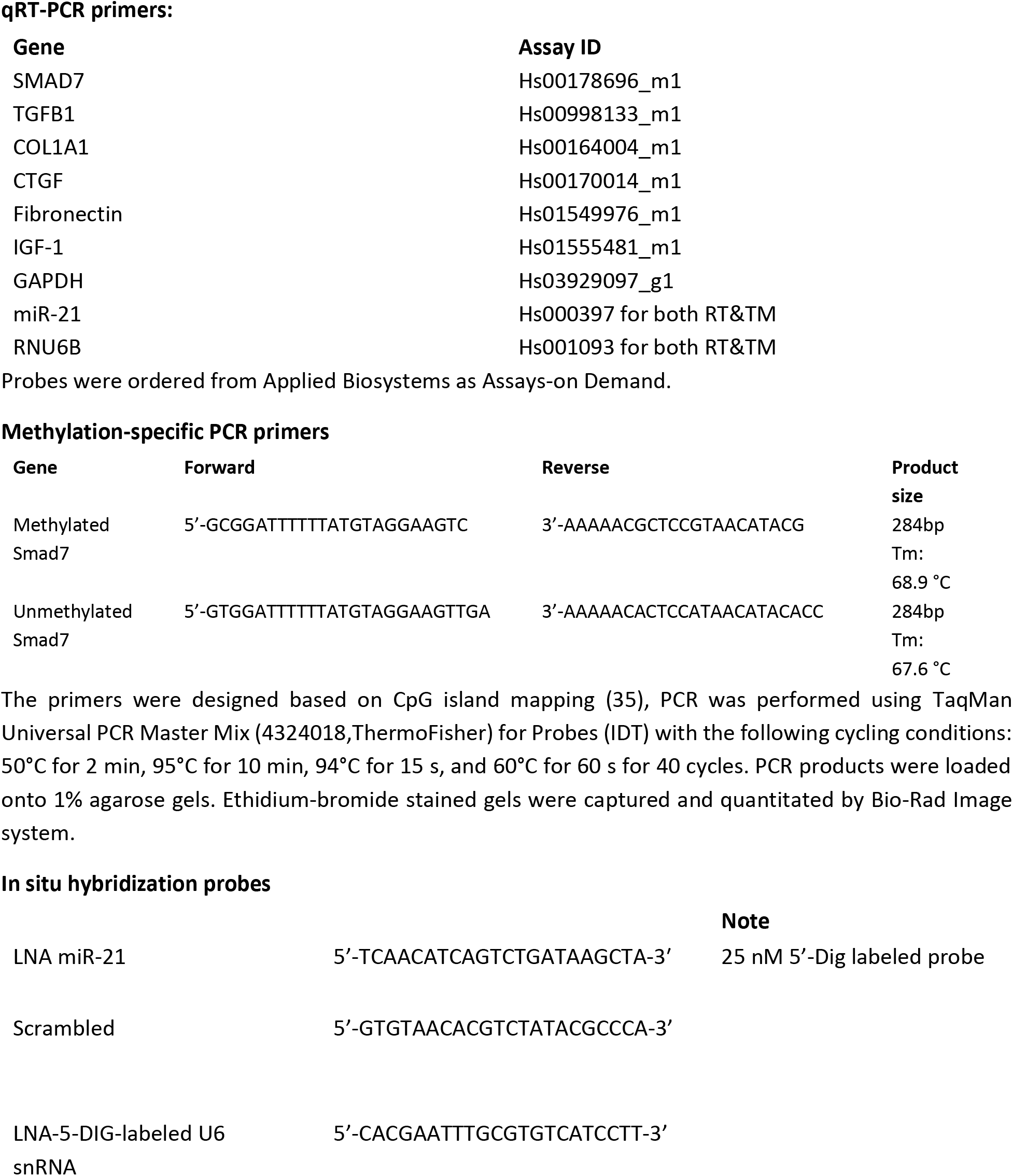
Subject demographics:

A unique feature of this study was that SEMF were isolated from both affected ileum and histologically normal proximal ileal resection margin (normal ileum) in the same patient with Crohn’s disease, as well as ileal tissue obtained from non-Crohn’s subjects undergoing surgery for nonmalignant conditions, including trauma ^35,36^.

Human intestinal SEMFs were isolated by enzymatic digestion as previously described ^36^. Briefly, ileal mucosa was removed by shaking at 37°C in Hank’s balanced salt solution containing 5 mM EDTA. Samples denuded of epithelial cells were rinsed with ice-cold PBS and then incubated in 20 ml of RPMI-5 medium supplemented with 10% fetal bovine serum, 2 mM L-glutamine, 10mM HEPES, pH7.4, 1mM sodium pyruvate, 100 U/mL penicillin, 100 µg/mL streptomycin, 20 U of dispase II, and 4,000 U collagenase D (Roche, 11088858001). Tissue was shaken vertically at 250 rpm for 1 hour at 37°C. The resultant cell suspension was centrifuged at 200 g for 5 minutes at 4°C, washed once, resuspended in 10 ml RPMI-5 complete medium, and cultured at 37°C in a humidified incubator containing 5% CO2. Nonadherent cells were removed after every 24h. SEMFs were subsequently used for RNA and microRNA isolation, whole cell lysates preparation, or primary cell culture experiments, as previously reported and validated ^5,35,36^.

Cells isolated using this protocol are devoid of epithelial cells, endothelial cells, neurons, and interstitial cells of Cajal. Instead, they exhibit a myofibroblast phenotype characterized by positive immunostaining for myofibroblast markers and negative staining for fibroblast markers ^35,37^. These phenotypic characteristics, as well as the epigenetic marks observed in vivo, are retained during primary culture ^35,37^.

### Ethical Considerations

All human studies were approved by the Virginia Commonwealth University Institutional Review Board. Written informed consent was obtained from all participants prior to specimen collection.

### Quantitative real-time PCR

Quantitative real-time PCR (qRT-PCR) was used to quantify RNA transcript expression. Relative expression levels were calculated using the 2^-ΔΔCt^ method, with glyceraldehyde-3-phosphate dehydrogenase (GAPDH) used as the endogenous control for mRNA analysis and U6 used for miRNA analysis ^38,39^. Primer sequences are listed in Supplementary Material, Table 2, and were purchased from Applied Biosystems (Foster City, CA). Methylated and unmethylated Smad7 expressions were also assessed using semiquantitative PCR. For methylation analysis of the human Smad7 promoter, primer sets specific for methylated and unmethylated Smad7 sequences were used (Supplementary Material, Table 2).

### *In situ* hybridization of hsa-miR-21

In situ hybridization for miR-21 was performed using the miRCURY LNA microRNA ISH Optimization Kit (Exiqon, 90002) together with 5′- and 3′-DIG-labeled probes specific to hsa-miR-21, according to the manufacturer’s instructions. The sequences of the LNA miR-21 probe, scrambled control probe, and LNA-5-DIG-labeled U6 snRNA probe are provided in Supplementary Material, Table 2.

### RNAscope

SEMFs were seeded into 8-well chamber slides at a density of 5 × 10^4^ cells/well and were either left untreated or stimulated with TGF-β1 (10 ng/mL) for 16 hours. Cells were fixed in prechilled 10% neutral-buffered formalin for 30 minutes at room temperature. Slides were dehydrated sequentially in 50%, 70%, and 100% ethanol for 5 minutes each, followed by rehydration in 70% and 50% ethanol for 2 minutes each at room temperature.

Slides were treated with RNAscope® Hydrogen Peroxide for 10 minutes and subsequently incubated with RNAscope® Protease III for 10 minutes at room temperature. Target probes for pre-miR-21, a negative control, and a positive control were then hybridized for 2 hours at 40°C. Signal amplification was performed according to the manufacturer’s instructions using the BaseScope™ Detection Reagent v2-Red Assay (Advanced Cell Diagnostics, Hayward, CA).

Amplification reagents AMP1, AMP2, AMP3, AMP4, AMP5, and AMP6 were applied sequentially for 30, 30, 15, 30, 30, and 15 minutes, respectively, at 40°C. AMP7 and AMP8 were subsequently applied for 30 and 15 minutes, respectively, at room temperature. Before each amplification step, slides were washed twice with 1× wash buffer (Cat. No. 310091). Samples were then incubated with RED substrate solution for 10 minutes at room temperature protected from light.

Slides were rinsed with tap water and counterstained with 50% Gill’s Hematoxylin I (American MasterTech Scientific) for 2 minutes at room temperature. Following rinsing, slides were immersed in 0.02% ammonia water and washed again with tap water. Samples were completely air-dried, mounted with VectaMount mounting medium, and coverslipped. Images were acquired using an Olympus brightfield microscope.

For quantification, 20 images per sample were collected from five regions of interest (four quadrants and one central region). Four random fields were captured at 20× magnification within each region. Pre-miR-21 RNAscope signals were quantified using Fiji/ImageJ software. For chromogenic assays, images were acquired using a standard brightfield microscope equipped with a digital camera at 20× magnification. Results were expressed as the average number of red punctate signals per cell ^40^.

RNAscope staining was scored semi-quantitatively using a five-tier grading system:

| Score | Criteria (20× Magnification) |
|---|---|
| 0 | No staining or fewer than 1 dot per 10 cells |
| 1+ | 1-3 dots per cell |
| 2+ | 4-10 dots per cell; very few dot clusters |
| 3+ | >10 dots per cell; <10% of positive cells contain dot clusters |
| 4+ | >10 dots per cell; >10% of positive cells contain dot clusters |

### Immunoblot Analysis

Cell lysates were prepared as previously described ^41–43^. Protein expression levels of phospho-Smad3 (Ser423/425), total Smad3 (Cell Signaling Technology, Danvers, MA), collagen I, CTGF, IGF-I, fibronectin (FN), Smurf1, Smurf2, HDAC1, p300, ubiquitin (Santa Cruz Biotechnology, Santa Cruz, CA), and acetyl-lysine (Cytoskeleton, Inc., Denver, CO) were determined by immunoblot analysis and normalized to β-actin (Sigma-Aldrich, St. Louis, MO) ^41,44^.

### RNA-Chromatin immunoprecipitation (RNA-ChIP)

RNA-ChIP was performed on nuclear extracts from freshly isolated cells. Cells were used for transfection with Smad3 mutants and treatment with TGF-β1 (10 ng/ml) for 16 hours prior to extraction of genomic DNA using TRIzol reagent (Invitrogen, Carlsbad, CA) and quantitative analysis of Smad3 binding activity to pri-miR21. ChIP was conducted using a ChIP-grade anti-Smad3 antibody (Cell Signaling Technology, Danvers, MA) or control rabbit IgG. PCR was performed using primers specific to the pri-miR-21 promoter region (SwitchGear Genomics, Menlo Park, CA). Results were calculated from input, PCR performed with DNA isolated from nuclear extracts without immunoprecipitation. Binding activity was quantified using the following formula:

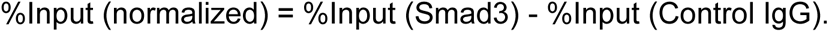

### Luciferase Reporter Assay

SEMFs were transfected with 1 μg of a dual-reporter construct for pri-miR21 (HPRM13178-PG04, GeneCopoeia, Rockville MD) using EndoFectin™ Lenti transfection reagent (EFP1003, GeneCopoeia, Rockville MD). After 16 hours of transfection cells were treated with IL-6 (10 ng/ml) for 6 hours. Medium was collected for Gaussia Luciferase and secreted alkaline phosphatase luminescence assays. Luminescence assays were performed in triplicate using a Wallac Victor2 1420 Multilabel counter (Perkin Elmer Life Sciences, Waltham, MA).

### Transfection of anti-hsa-miR-21 and overexpression of pre-hsa-miR-21

Transfection was performed using X-tremeGENE siRNA (Roche) transfection reagent using 5’-FAM-tagged oligonucleotides: anti-hsa-miR-21 (AntagomiR-21), pre-hsa-miR-21, or scrambled controls (Exiqon). For each transfection, 100 nM of anti-miR or pre-miR was diluted in 250 μl Opti-MEM (Lonza). Total RNA was subsequently extracted using TRIzol reagent. Transfection efficiency exceeded 75%, as determined by fluorescence microscopy. When the precursor pre-hsa-miR-21 is overexpressed, it undergoes enzymatic cleavage by Dicer to yield active mature strands (hsa-miR-21-5p and hsa-miR-21-3p).

### Manipulation of Smad3 function

Smad3 activity was manipulated using three approaches: siRNA-mediated knockdown, expression of a dominant negative (dn)Smad3(3ΔC) mutant, and overexpression of wildtype Smad3 that functions as constitutively active Smad3 ^45^. For siRNA-mediated knockdown, SEMFs were transfected with 20 nM Smad3 siRNA or scrambled control siRNA using Opti-MEM I Reduced Serum Medium (Thermo Fisher Scientific, Waltham, MA). For overexpression studies, 1 μg of plasmid containing either wild type Smad3 or (dn)Smad3(3ΔC) mutant was transfected into primary SEMFs and then incubated for 24-48 hours before RNA extraction using TRIzol reagent (Invitrogen, Carlsbad, CA).

### Overexpression of Smad7

Smad7 expression was enhanced using pCMV5-Smad7-HA and pBabe Flag Smad7 plasmids. The pCMV5-Smad7-HA was a gift received from Jeff Wrana (Addgene plasmid # 11733; http://n2t.net/addgene:11733; RRID:Addgene_11733), whereas pBabe Flag Smad7 was a gift received from Rik Derynck (Addgene plasmid # 14836; http://n2t.net/addgene:14836; RRID:Addgene_14836).

For TGFβ-inducible luciferase reporter assays, HepG2 cells were transiently transfected with the reporter plasmid (p3TP-Lux), pCMVβ-gal and the indicated constructs or with empty vector alone. To induce the luciferase reporter, cells were treated with TGF-β1 (10 ng/ml).

### Confocal microscopy

Human intestinal tissue cryosections (7 μm thick) were used for microscopic analysis. Following in situ hybridization, pre-hsa-miR-21 localization was visualized by light microscopy as previously described ^3,46^. Digital images were acquired using a Leica TCS-SP2 AOBS confocal laser-scanning microscope.

### Statistical analysis

Data are presented as mean ± SEM. The value of n represents the number of independent experiments performed using tissues or cells from different subjects.

Comparisons between two groups were performed using paired or unpaired Student’s t-tests, as appropriate. Comparisons among three or more groups were conducted using one-way analysis of variance (ANOVA) followed by Tukey’s multiple-comparison post hoc test. Statistical significance was defined as P < 0.05.

## Results

### Smad7 expression is diminished in mesenchymal cells of the affected intestine

Our previous work identified sustained, uninhibited activation of canonical TGF-β1 signaling through Smad3 in mesenchymal cells isolated from the affected ileum of patients with Montreal B2 fibrostenotic phenotype of CD ^3^. To investigate why Smad3-dependent TGF-β1 signaling remained persistently activated and failed to respond to normal negative feedback, we examined expression of Smad7, a key inhibitor of canonical TGF-β1 signaling. Smad7 expression was measured in SEMFs isolated from affected and uninvolved ileal regions of patients with Montreal B1, B2, or B3 CD and compared with SEMFs from non-Crohn’s control subjects. Smad7 transcript levels were increased in affected ileum from patients with B1 inflammatory disease compared with their matched uninvolved ileum and with non-Crohn’s controls (Figure 1A). Smad7 expression was unchanged in patients with B3 penetrating disease. These findings are consistent with previous reports showing increased Smad7 protein expression in mucosal biopsy specimens from patients with CD ^47^.

**Figure 1:**
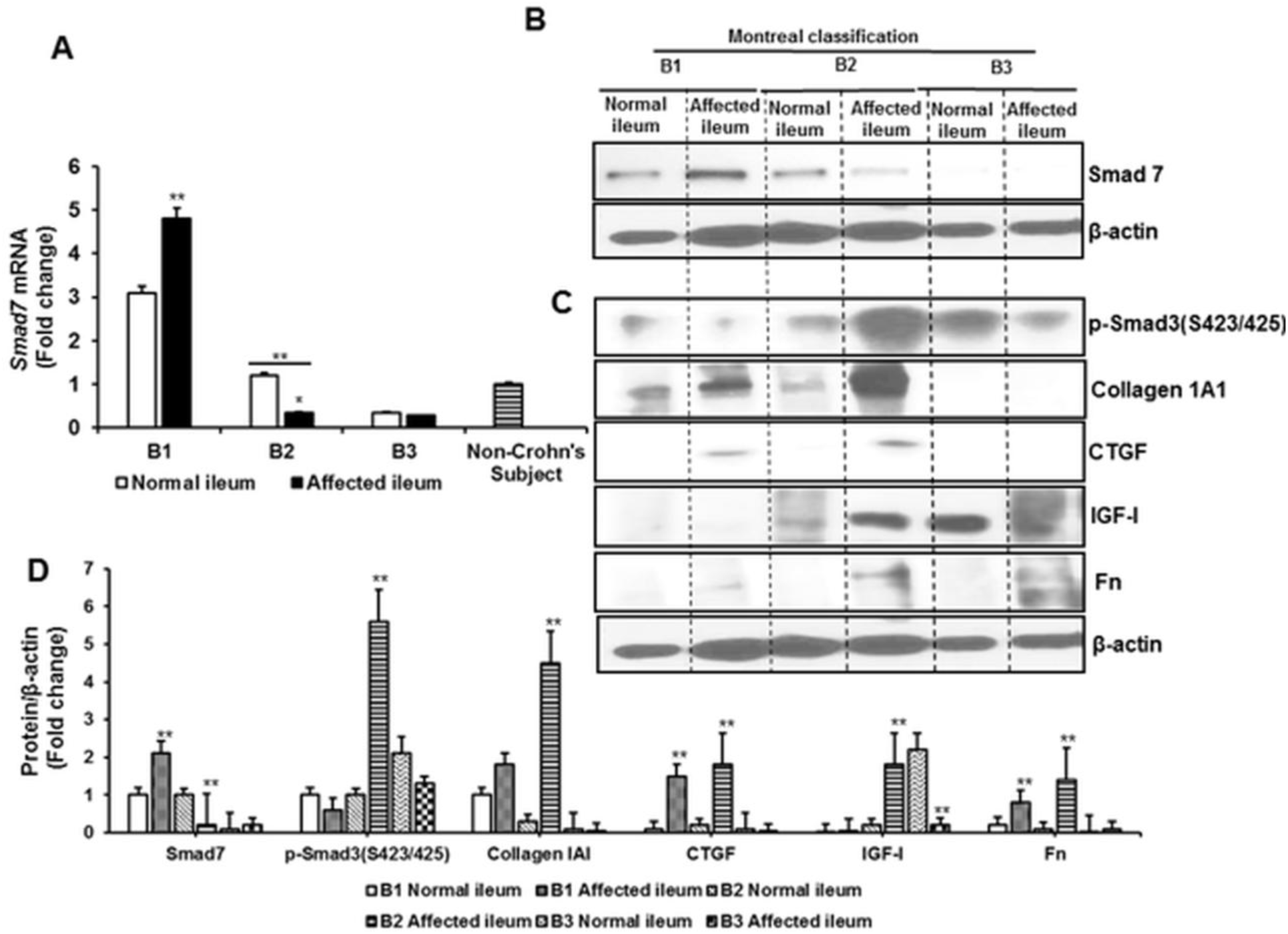
Distinct patterns of Smad7 and TGF-β1-dependent proteins in normal subjects and in each CD phenotype. *(A)* Smad7 transcript levels in affected ileum from the patients with B2 fibrostenotic phenotype are decreased compared to normal ileum in the same patient. Different patterns were normal ileum and affected ileum in each CD phenotype. *(B)* Representative immunoblot of Smad7 protein levels in normal ileum and affected ileum in the same subject in each CD phenotype. *(C)* Representative immunoblot of TGF-β1-dependent protein levels in normal ileum and affected ileum of the same subject in each CD phenotype. (D) Quantitative analysis of immunoblots in panels B and D showing a distinct pattern or decreased Smad7 and increased TGF-β1-dependent proteins levels in the affected ileum of B2 phenotype CD subjects compared to normal ileum in the same subject and compared to other CD phenotypes. Values are expressed as fold change in mRNA transcripts or protein levels and reported as mean ± SEM of 4-6 experiments from different subjects. * denotes p < 0.05 vs non-Crohn’s subjects, ** denotes P < 0.05 vs normal ileum in the same subject. Each group is representative of 6 subjects with the distinct phenotype indicated in the figure.

In contrast, Smad7 transcript levels were significantly decreased in affected ileum from patients with B2 fibrostenotic phenotype compared with uninvolved ileum from the same patient and with non-Crohn’s controls (Figure 1A). This reduction in Smad7 mRNA was accompanied by a corresponding decrease in Smad7 protein expression despite sustained TGF-β1 signaling and persistent Smad3 phosphorylation (Figure 1B-D) ^3^.

We next examined the consequences of reduced Smad7 expression and increased TGF-β1 signaling on Smad3 activity and the expression of collagen Iα1 and other fibrosis-associated proteins. Collagen Iα1 protein levels were significantly increased in affected ileum compared with uninvolved ileum from the same patient with B2 CD (Figure 1C,D). This increase was not observed in patients with B1 or B3 disease. Similarly, the TGF-β1-responsive profibrotic proteins connective tissue growth factor (CTGF), insulin-like growth factor I (IGF-I), and fibronectin (FN) were all elevated in fibrostenotic ileum compared with paired uninvolved ileum (Figure 1C,D).

Notably, the characteristic profile of increased TGF-β1, pSmad3(S423/425), collagen Iα1, CTGF, IGF-I, and FN expression accompanied by reduced Smad7 levels was observed in SEMF from all the patients with fibrostenotic phenotype included in this study, regardless of treatment with aminosalicylates, immunomodulators, or biologic therapies.

### hsa-miR-21 expression is increased in mesenchymal cells of the affected intestine

Microarray analyses have identified increased hsa-miR-21 expression in the heterogeneous cell populations present in endoscopic mucosal pinch biopsies from patients with Crohn’s disease. However, the abundance of hsa-miR-21 in ileal SEMFs and the mechanisms regulating its expression remain poorly defined ^18,48^.

To address this question, we first examined hsa-miR-21 expression in SEMFs by qRT-PCR. Expression levels of hsa-miR-21 were elevated in SEMFs isolated from normal ileum of patients with B2 fibrostenotic CD compared with non-Crohn’s controls and were further increased in SEMF from affected ileum (Figure 2A). Increased hsa-miR-21 expression was also observed in SEMF from patients with B1 inflammatory Crohn’s disease. In contrast, no significant change in hsa-miR-21 expression was detected in patients with Montreal B3 penetrating disease.

**Figure 2:**
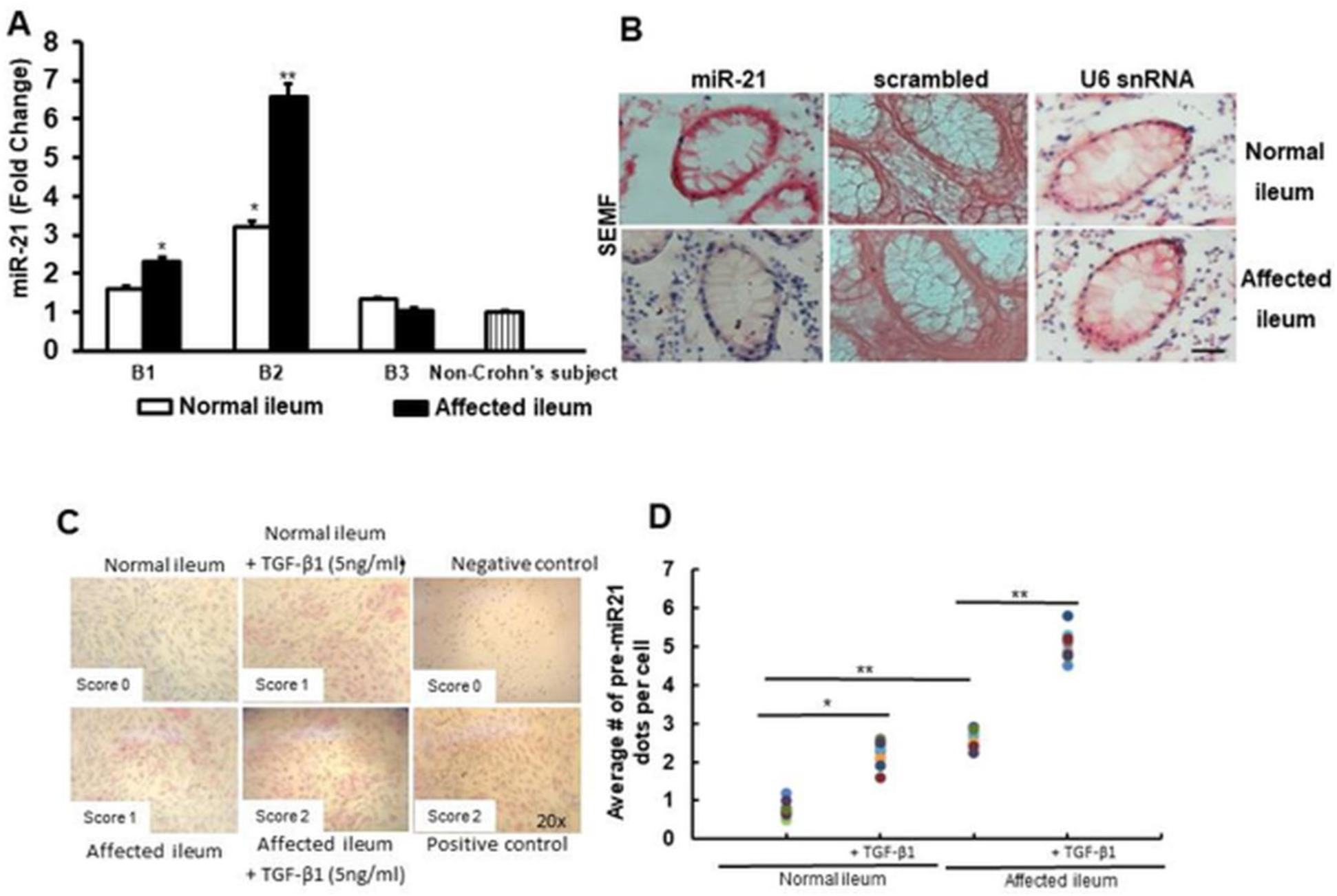
Expression and localization of hsa-miR-21 in patients with fibrostenotic Crohn’s disease. *(A)* hsa-miR-21 expression is increased in affected ileum compared to normal ileum in the same subjects with B2 fibrostenotic phenotype and compared to other CD phenotypes and to non-Crohn’s subjects. Values represent the mean ± SEM of 4-6 experiments using different subjects. Results were normalized to U6 snRNA and expressed as fold change from non-Crohn’s subjects. * denotes p < 0.05 vs non-Crohn’s subjects; ** denotes p < 0.05 vs normal ileum region in the same subject. (B) Representative *in situ* hybridization of hsa-miR-21 in 4 separate Montreal B2 subjects’ histologic sections at 20x magnification. U6 snRNA served as positive control. Hsa-miR-21 is shown in blue with red nuclear counterstain. (C) Representative RNAScope *in situ* hybridization analysis of pre-miR21 in primary cell culture of SEMF isolated from normal ileum and affected ileum in the same subject with B2 fibrostenotic CD phenotype. Cells were treated with TGF-β1(10 ng/ml) for 16 hours and subjected to RNAScope in situ hybridization using pre-miR21, negative control, and positive control probes. Images were captured at 20x magnification. (D) Quantitative analysis of positive red dots of pre-miR21 RNA per cell under the indicated conditions. Scoring standard is referred to methods section for details. Black scale bar=150 µm. Each group data is representative of 6 subjects with a Montreal B2 CD phenotype indicate. * denotes p < 0.05 vs normal ileum. ** denotes p < 0.05 between normal ileum vs affected ileum, or affected ileum vs affected ileum without TGF-β1 treatment.

These observations were further validated using *in situ* hybridization and RNAscope analyses of histological sections from patients with B2 Crohn’s disease. *In situ* hybridization demonstrated increased hsa-miR-21 expression specifically within SEMFs, whereas epithelial cells showed no appreciable change (Figure 2B). This increase was observed in affected ileum relative to paired uninvolved ileum from the same patient. The findings were corroborated by RNAscope analysis of pre-hsa-miR-21 levels in cultured SEMFs (Figure 2C). In untreated SEMF isolated from normal ileum, pre-hsa-miR-21 levels were low and similar to those observed in negative control samples. In contrast, untreated SEMF from fibrostenotic ileum exhibited substantially higher levels of pre-hsa-miR-21. Treatment with TGF-β1 further increased pre-hsa-miR-21 expression in both normal and fibrostenotic SEMF (Figure 2D).

### hsa-miR-21 regulates Smad7 mRNA expression

Hybridization analysis identified an 8-mer consensus seed sequence complementary to hsa-miR-21-5p within nucleotides 1122-1129 of the SMAD7 3′-UTR, suggesting that miR-21 may directly regulate Smad7 mRNA abundance (Figure 3A) ^49^. We therefore examined whether hsa-miR-21 regulates Smad7 expression and, consequently, TGF-β1 signaling and the expression of TGF-β1-responsive profibrotic genes, including *COL1A1, IGFI, FN,* and *CTGF*.

**Figure 3:**
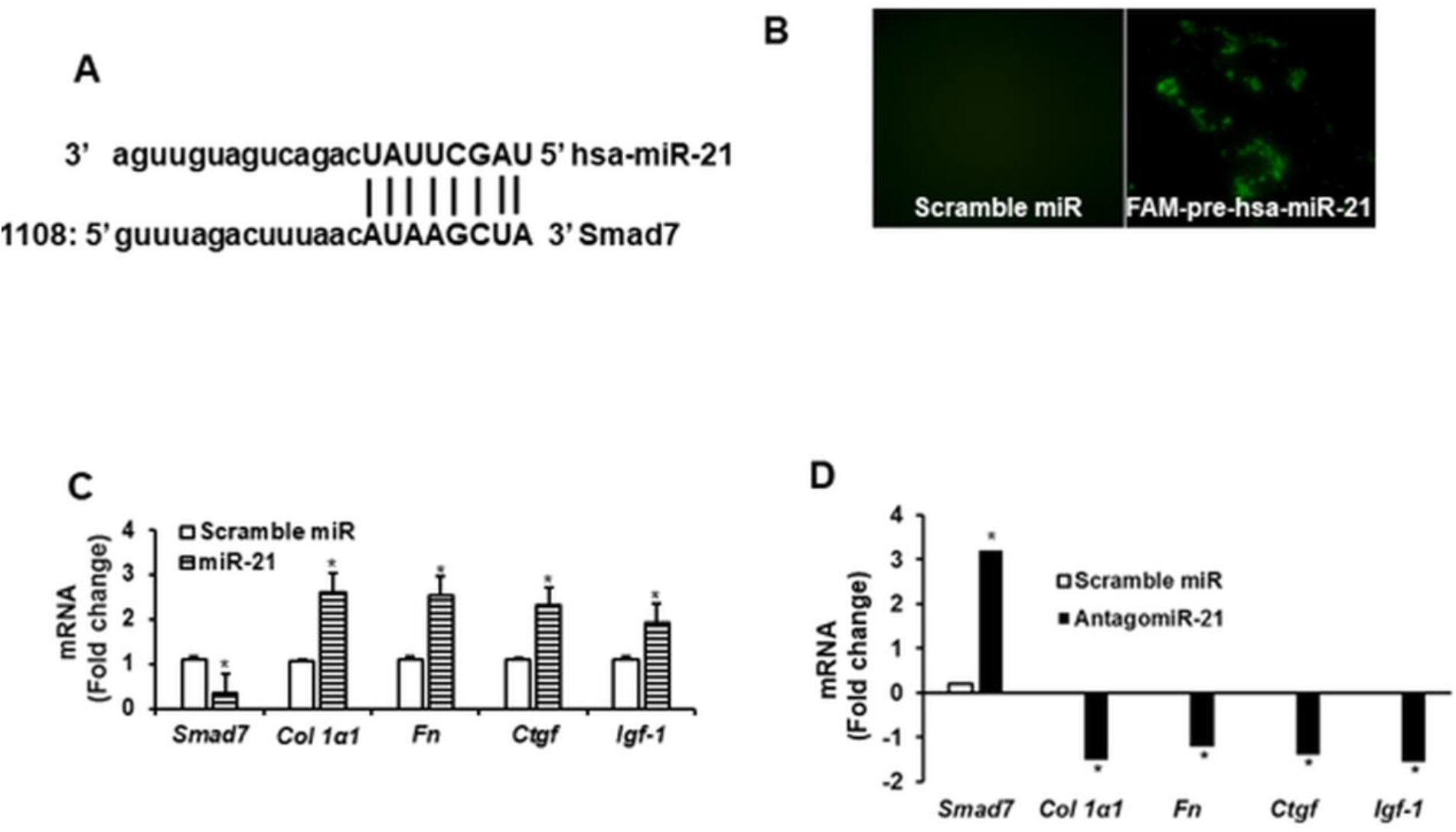
miR-21 directly targets Smad7 and affects profibrotic genes expression. *(A)* The human Smad7 3’-UTR fragment alignment with the 8-mer hsa-miR-21 sequence. *(B)* Upper panel: Representative image of FAM-miR-21 transfection in SEMF (>80% efficiency). *(C)*: miR-21 transfection decreased Smad7 expression and increased TGF-β1-dependent gene expression compared to scrambled sense miR control. *(D)* Transfection of FAM-antagomiR-21 in cultured SEMF from affected ileum in patients with B2 fibrostenotic phenotype increased Smad7 and decreased TGF-β1-dependent gene expression compared to the scramble sense antagomiR control. Results were expressed as mean ± SEM of 4-6 experiments on SEMF isolated from different subjects and reported as fold change from control. * denotes p < 0.05 vs scrambled sense antagomiR control.

Transfection of FAM labeled pre-hsa-miR-21 into SEMFs isolated from normal ileum of B2 patients achieved greater than 80% transfection efficiency (Figure 3B). Overexpression of hsa-miR-21 reduced Smad7 expression and was accompanied by increased expression of *TGFB1, COL1A1, FN, CTGF,* and *IGFI* (Figure 3C).

Conversely, transfection of antagomiR-21 into SEMFs isolated from fibrostenotic ileum produced the opposite effect. Smad7 expression increased, whereas expression of *TGFB1, COL1A1, FN, CTGF,* and *IGFI* decreased (Figure 3D). These findings indicate that hsa-miR-21 negatively regulates Smad7 expression and promotes activation of a profibrotic transcriptional program in intestinal mesenchymal cells.

### Smad3 regulates pri-hsa-miR-21 transcription

We next investigated the mechanisms responsible for increased hsa-miR-21 expression in SEMF. Importantly, intestinal mesenchymal cells retain key features of their fibrotic phenotype in primary culture, including epigenetic modifications and post-translational regulatory mechanisms^35,38^. This property allowed us to determine whether persistent Smad3(S423/425) phosphorylation contributes to elevated hsa-miR-21 expression.

Smad proteins are known to bind conserved RNA motifs and facilitate Drosha-mediated processing of precursor miRNAs ^21^. However, the pri-hsa-miR-21 promoter contains three consensus Smad-binding elements (5′-CAGAC-3′), suggesting that Smad3 may directly regulate pri-hsa-miR-21 transcription and not solely via Drosha-dependent miRNA maturation ^21,22^. To test this possibility, Smad3 binding to the pri-hsa-miR-21 promoter was assessed by RNA-ChIP. Basal Smad3 binding to the pri-hsa-miR-21 promoter was substantially greater in SEMFs from fibrostenotic ileum than in SEMFs from paired uninvolved ileum. Treatment with SIS3, an inhibitor of Smad3(S423/425) phosphorylation, markedly reduced Smad3 binding to the pri-hsa-miR-21 promoter (Figure 4A) ^50^.

**Figure 4:**
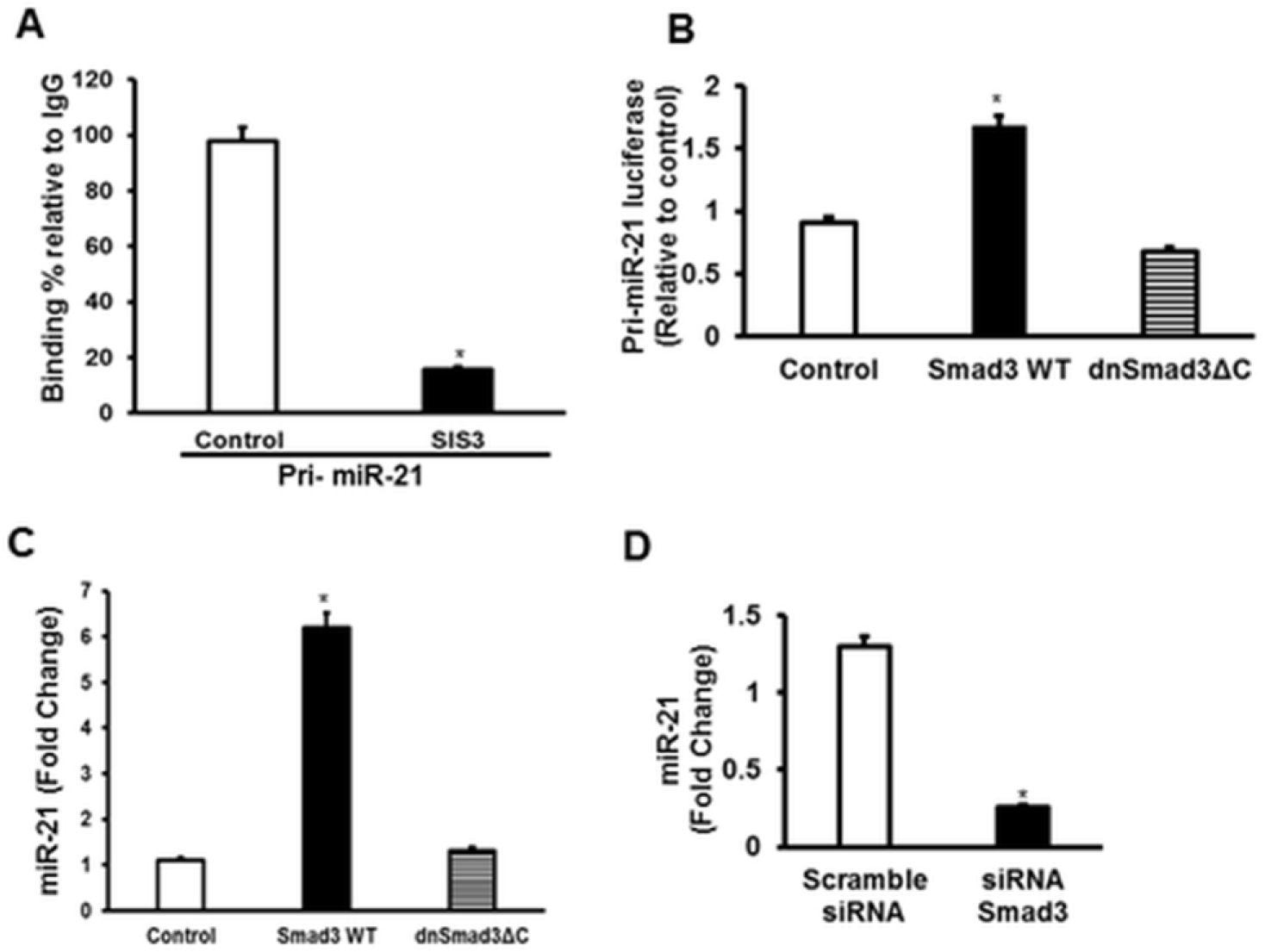
Smad3 regulated pre-miR-21 transcription increases miR-21 expression. *(A)* Binding affinity of phosphorylated Smad3 to the pri-miR-21 promoter in SEMFs isolated from affected ileum was inhibited by SIS3, an inhibitor of Smad3 phosphorylation. *(B)* Transcriptional activity of pri-miR-21 was increased by transfection of wild type Smad3 (WT) (mimicking constitutively active Smad3) but not affected by transfection of dnSmad3ΔC. (C) miR-21expression was increased in SEMF transfected with wtSmad3 compared to empty vector. miR-21 expression was not affected by transfection of a dnSmad3ΔC. *(D)* RNAi mediated knockdown of Smad3 in SEMF decreased miR-21 expression compared to scrambled sense RNAi. Values represent the mean ± SEM of 4-6 experiments using SEMF isolated from different subjects with B2 fibrostenotic CD and reported as fold change from control. * denotes p < 0.05 vs control or scrambled sense siRNA.

The ability of Smad3 to regulate pri-hsa-miR-21 transcription was further confirmed using luciferase reporter assay and qRT-PCR. Overexpression of wild-type Smad3, which functions as a constitutively active form of Smad3, increased pri-hsa-miR-21 transcription activity (Figure 4B) and enhanced expression of mature hsa-miR-21 transcripts (Figure 4C). In contrast, expression of a dominant-negative Smad3 mutant (dn)Smad3(3ΔC) did not significantly suppress pri-hsa-miR-21 transcription or hsa-miR-21 expression relative to empty vector control (Figure 4B,C). This likely reflects the low basal level of Smad3 activation present in SEMFs derived from normal ileum, even in patients with fibrostenotic CD (Figure 1B,D) ^51^. Collectively, these findings support a model in which activated Smad3 directly regulates pri-hsa-miR-21 transcription in SEMFs from fibrostenotic ileum.

### Post-translational acetylation of Smad7

Smad7 has been identified as a direct target of the transcriptional coactivator p300 both *in vitro* and *in vivo* ^32^.The stability of Smad7 is regulated by the balance between p300 acetyltransferase and histone deacetylase activities. Acetylation of Smad7 at K70 and K74 protects the protein from Smurf1/2 E3 ubiquitin ligase -dependent polyubiquitination and subsequent 26S proteasomal degradation, whereas deacetylation promotes ubiquitination and protein degradation ^32–34^. Thus, acetylation of these residues stabilizes Smad7 protein and protects it from Smurf-dependent degradation ^32^.

Here we showed both protein levels of p300 and histone deacetylase 1 (HDAC1) were elevated in SEMFs isolated from affected ileum of patients with B2 fibrostenotic CD compared with matched normal ileum (Figure 5A). However, quantitative analysis of protein bands from Western blots revealed that the increase in HDAC1 protein level was significantly greater than that in p300 expression (Figure 5B). Importantly, these alterations were not observed in patients with B1 or B3 disease, in whom Smad7 expression remained appropriately regulated (Figure 1A,B).

**Figure 5.**
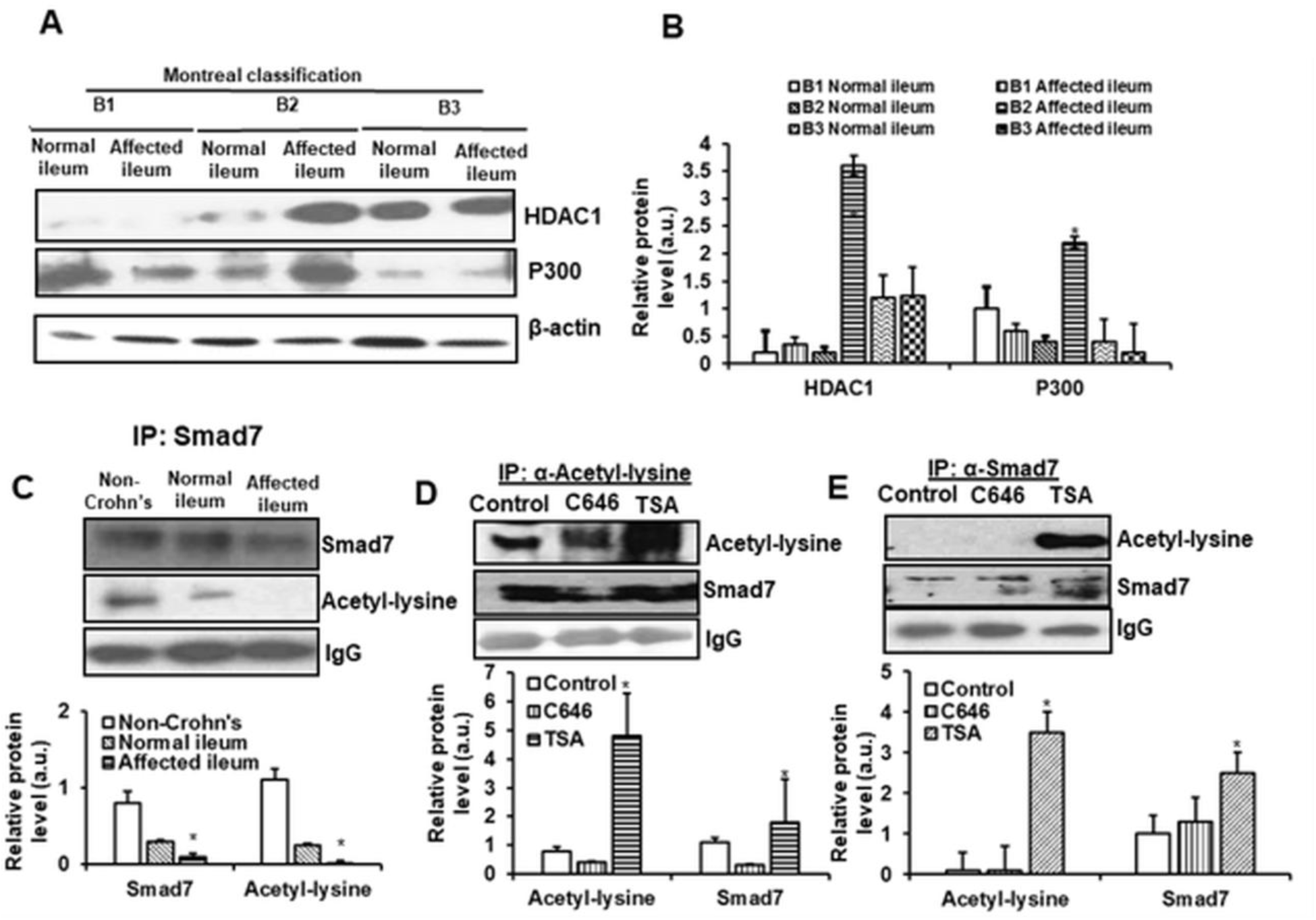
Lysine acetylation regulates Smad7 levels. *(A)* Representative immunoblots (left panel) and (B) densitometric analysis (right panel) showing increased HDAC1 and p300/CBP acetyltransferase levels in affected ileum of fibrostenotic B2 Crohn’s disease subjects compared to normal ileum in the same subject and compared to other CD phenotypes. (C) Representative co-immunoprecipitation (upper panel) and densitometric analysis (lower panel) of lysine acetylation of Smad7 and Smad7 levels that are decreased in SEMF of normal ileum of subjects with B2 fibrostenotic CD that is further decreased in the affected ileum in the same patient. *(D* & *(E)* Representative co-immunoprecipitation (upper panel) immunoblots and quantitative densitometry (lower panel) showing lysine acetylation of Smad7 and Smad7 levels in SEMF of fibrostenotic ileum that are decreased by the p300/CBP acetylation inhibitor, C646 and increased by the histone deacetylase inhibitor, Trichostatin A. IgG or β-actin served as loading control for the IP/IB experiments. Results represent the mean ± SEM of 4-6 experiments using different subjects’ samples. Values are expressed in arbitrary units (a.u.). * denotes p < 0.05 vs non-Crohn’s or normalized to non-treated control. Each group data is representative of 6 patients with distinctive phenotype indicated in the figure.

These findings suggested that overall net Smad7 deacetylation is favored in SEMFs from affected ileum of patients with B2 fibrostenotic Crohn’s disease. Consistent with this hypothesis, the abundance of acetylated Smad7 was significantly reduced in SEMFs from affected ileum compared with SEMFs from non-Crohn’s controls and normal ileum from patients with B2 fibrostenotic CD (Figure 5C).

To confirm that acetylation of Smad7 on lysine amino acid directly influences protein stability, SEMFs from normal ileum of patients with B2 fibrostenotic CD were treated with either a p300 inhibitor, C646, or a HDAC1 inhibitor, Trichostatin A (TSA). Inhibition of p300 reduced acetylated Smad7 levels, whereas inhibition of HDAC1 significantly increased Smad7 acetylation, as demonstrated by immunoprecipitation analysis (Figure 5D,E) ^52,53^. Following immunoprecipitation, Smad7 and acetylated lysine residues were detected using anti-Smad7 and anti-acetyl-lysine antibodies, respectively (Figure 5D,E).

Together, these data demonstrate that reduced Smad7 acetylation contributes to diminished Smad7 stability in fibrostenotic CD and provide mechanistic evidence that altered p300-HDAC1 balance is an important mechanism regulating Smad7 protein abundance.

### SMURF1/2 mediates Smad7 ubiquitination

The susceptibility of Smad7 to Smurf-mediated E3 ubiquitination and subsequent proteasomal degradation is influenced by its acetylation status ^7,32,33^. We therefore examined the effects of TGF-β1 on the expression and intracellular localization of Smurf1 and Smurf2 in SEMF (Figure 6A).

**Figure 6.**
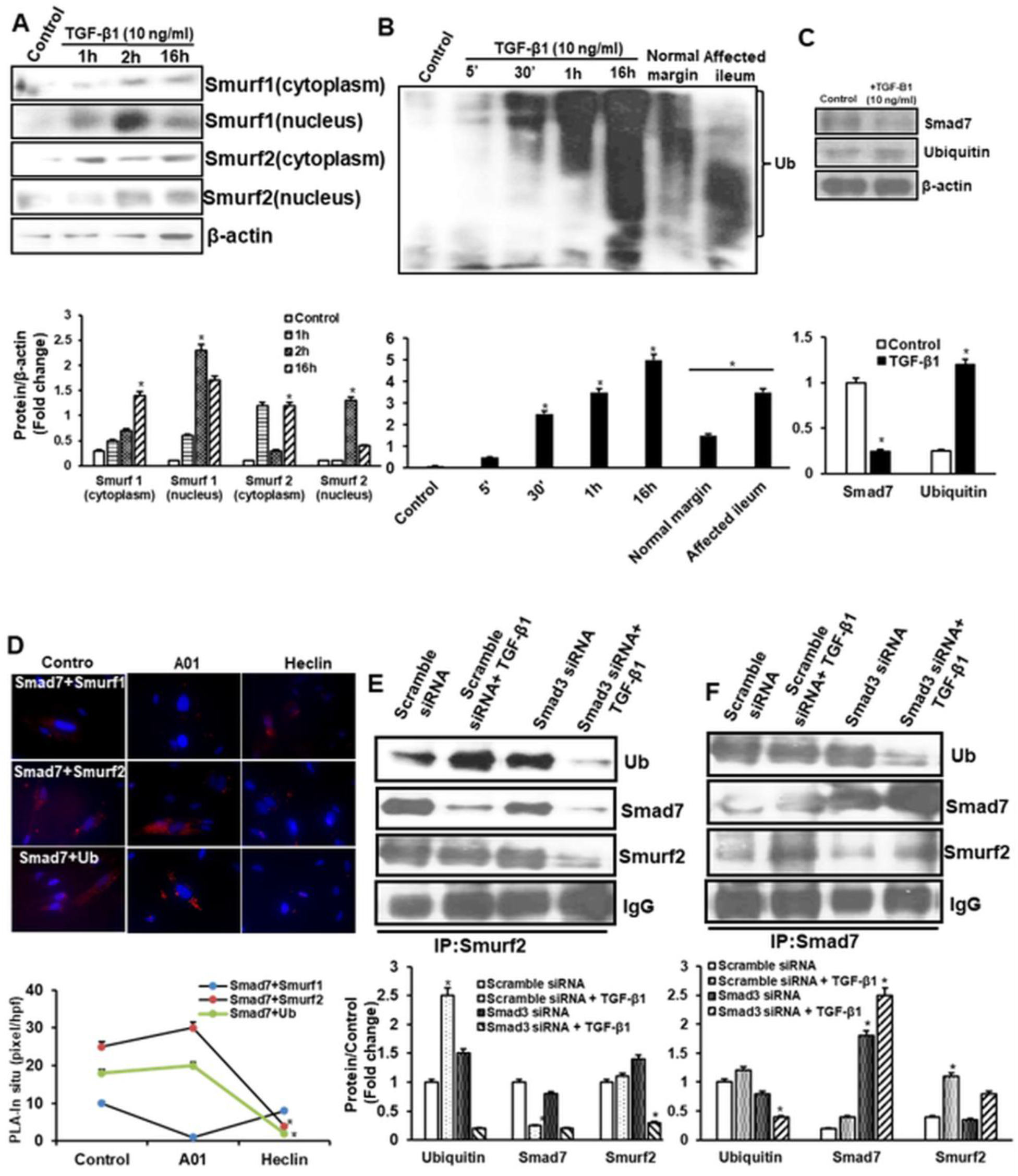
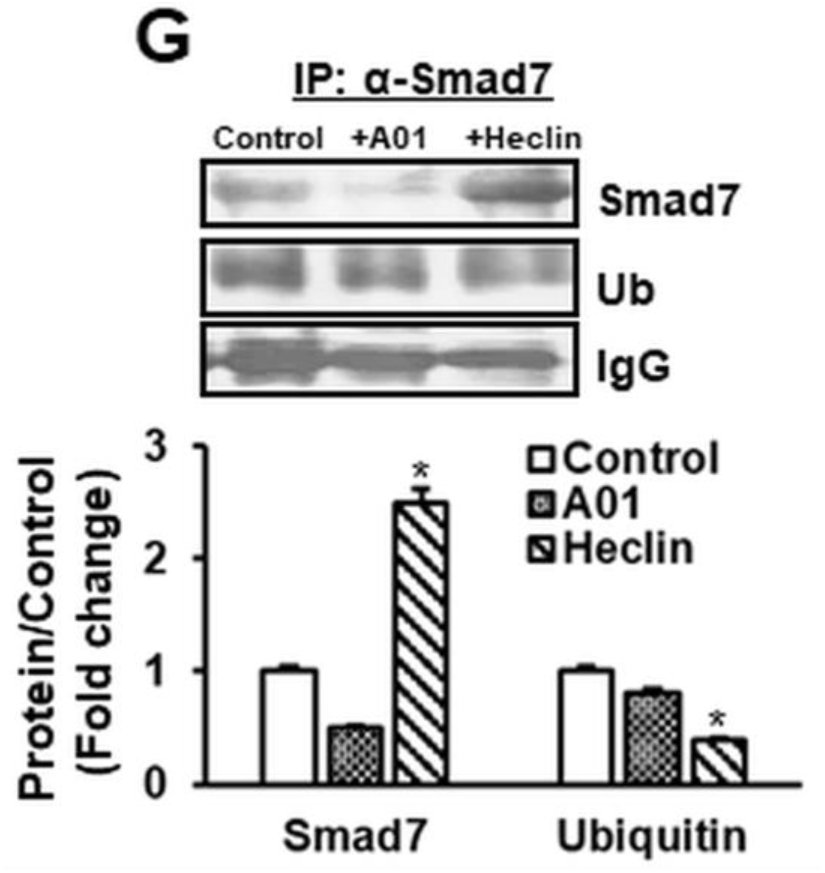
Smurf1/2-mediated ubiquitin-dependent degradation of Smad7. *(A)* Representative immunoblot and quantitative analysis of time-dependent nucleo-cytoplasmic translocation of Smurf1 and 2 proteins in SEMF of normal ileum in subjects with B2 fibrostenotic CD response to TGF-β1 (10 ng/ml). *(B)* Representative immunoblot and densitometric analysis of time-dependent ubiquitination of proteins in untreated SEMF of fibrostenotic ileum is increased compared to normal ileum in the same patient and mimics the response to TGF-β1 treatment. *(C)* Representative immunoblot and densitometric analysis of increased ubiquitination and decreased Smad7 levels in SEMF of normal ileum from subjects with fibrostenotic B2 CD in response to 16 h of TGF-β1 treatment. *(D)* Representative PLA imaging and PLA quantification of Smad7 interaction with Smurf1, Smurf2 and ubiquitin in SEMF from fibrostenotic ileum in the presence and absence of the Smurf1 inhibitor, A01 (µM) and the Smurf2 inhibitor, Heclin (µM). *(E)* Representative immunoblot and densitometric analysis of ubiquitin, Smad7, and Smurf2 after immunoprecipitation of Smurf2 in SEMF treated with TGF-β1 (10ng/ml) after siRNA-mediated knockdown of Smad3. *(F)* Representative immunoblot and densitometric analysis of ubiquitin, Smad7, and Smurf2 after immunoprecipitation of Smad7 in SEMF treated with TGF-β1 (10ng/ml) after siRNA-mediated knockdown of Smad3. *(G)* Representative immunoblot of Smad7 and ubiquitin after immunoprecipitation of Smad7 in SEMF of fibrostenotic ileum treated with A01 and heclin. IgG or β-actin served as loading control for the IP/IB experiments. Results represent the mean ± SEM of 4-6 experiments using different subjects’ samples with fibrostenotic CD from either the normal ileum or affected ileum as noted.. Values are expressed in arbitrary units (a.u.). * denotes p < 0.05 vs control, vs with A01 in (*D)*.

SEMFs isolated from normal ileum of patients with B2 fibrostenotic CD were treated with 10 ng/mL TGF-β1 for increasing durations. Nuclear levels of both Smurf1 and Smurf2 increased progressively during the first 2 hours of treatment and subsequently declined, coinciding with a gradual accumulation of both proteins in the cytoplasmic compartment (Figure 6A). SEMFs isolated from affected ileum of patients with B2 fibrostenotic CD exhibited greater degradation of ubiquitinated proteins than SEMF from paired normal ileum of the same B2 patient (Figure 6B). Similarly, TGF-β1 treatment of SEMFs isolated from normal ileum of patients with B2 fibrostenotic CD induced a time-dependent increase in ubiquitinated proteins (Figure 6B), which was accompanied by a progressive reduction in Smad7 protein abundance (Figure 6C).

Previous studies have reported that following TGF-β1 stimulation and Smad3 activation, Smad7 translocates from the nucleus to the cytoplasm and then plasma membrane in association with Smurf1 or Smurf2, where Smurf proteins promote ubiquitination and degradation of TGF-β receptors ^7,32^. To investigate the direct interactions among Smad7, Smurf1, Smurf2, and ubiquitin, we performed proximity ligation assays to assess protein-protein interactions and their subcellular localization (Figure 6D).

Basal levels of Smad7-Smurf1, Smad7-Smurf2, and Smad7-ubiquitin interactions were low in SEMFs isolated from affected ileum patients with B2 fibrostenotic CD (Figure 6D, control condition). Treatment with the selective Smurf1 inhibitor, AO1, or the selective Smurf2 inhibitor, Heclin, effectively disrupted Smad7-Smurf1 and Smad7-Smurf2 interactions, respectively. Notably, inhibition of Smurf2 with Heclin produced a substantially greater reduction in Smad7-ubiquitin association than inhibition of Smurf1 with AO1 (Figure 6D), suggesting that Smurf2 plays a dominant role in Smad7 ubiquitination during TGF-β1 signaling.

Smurf2 has been reported to promote multiple mono-ubiquitination events on Smad3 and inhibit the formation of both homotrimeric Smad3 and heterotrimeric Smad3-Smad4 complexes ^33,54^. Because Smad3 activation also induces expression of inhibitory Smads, such as Smad7, as part of a negative-feedback regulation of TGF-β1 signaling. Therefore, we investigated whether TGF-β1-induced Smad3 activation affects protein levels of Smurf2, ubiquitin, and Smad7, as well as their interactions.

Using Smurf2 immunoprecipitation pull-down method, we found that TGF-β1 stimulation reduced the association between Smurf2 and Smad7 while increasing the association between Smurf2 and ubiquitin (Figure 6E, second column). These changes were consistent with enhanced Smad7 degradation through the ubiquitin-proteasome pathway. Importantly, siRNA-mediated Smad3 knockdown prevented the TGF-β1-induced increase in Smurf2 expression and resulted in preservation of Smad7 protein levels despite TGF-β1 stimulation (Figure 6E, 3rd and 4th column & 6F, 3rd and 4th column). Under these conditions, Smad7 ubiquitination and degradation were markedly reduced. These findings indicate that Smurf2-mediated ubiquitination and degradation of Smad7 is dependent on Smad3 signaling. Consistent with this conclusion, inhibition of Smurf2 with Heclin restored endogenous Smad7 protein levels more effectively than inhibition of Smurf1 with AO1 in SEMF isolated from affected ileum of patients with B2 fibrostenotic CD (Figure 6G). Collectively, these results suggest that Smad3 is required for efficient Smurf2-dependent ubiquitination and degradation of Smad7 in fibrostenotic Crohn’s disease.

### Promoter hypermethylation silences Smad7 transcription

To determine whether epigenetic mechanisms contribute to reduced Smad7 expression in fibrostenotic Crohn’s disease, we first examined expression of DNA methyltransferase 1 (DNMT1) in SEMF isolated from affected and normal ileum from each phenotype. DNMT1 expression was significantly increased in fibrostenotic ileum from patients with B2 disease compared with matched uninvolved ileum (Supplementary Figure 1).

Previous studies have identified SMAD7 as a target of DNMT1-mediated regulation (22). Therefore, we assessed SMAD7 promoter methylation using methylation-specific PCR. SEMFs isolated from affected ileum of patients with B2 fibrostenotic CD exhibited substantially higher levels of gene expression of methylated SMAD7 relative to unmethylated SMAD7 compared with SEMFs from paired normal ileum (Figure 7A). In contrast, patients with B1 inflammatory and B3 penetrating disease exhibited the opposite pattern, with relatively greater levels of unmethylated SMAD7 (Figure 7A).

**Figure 7.**
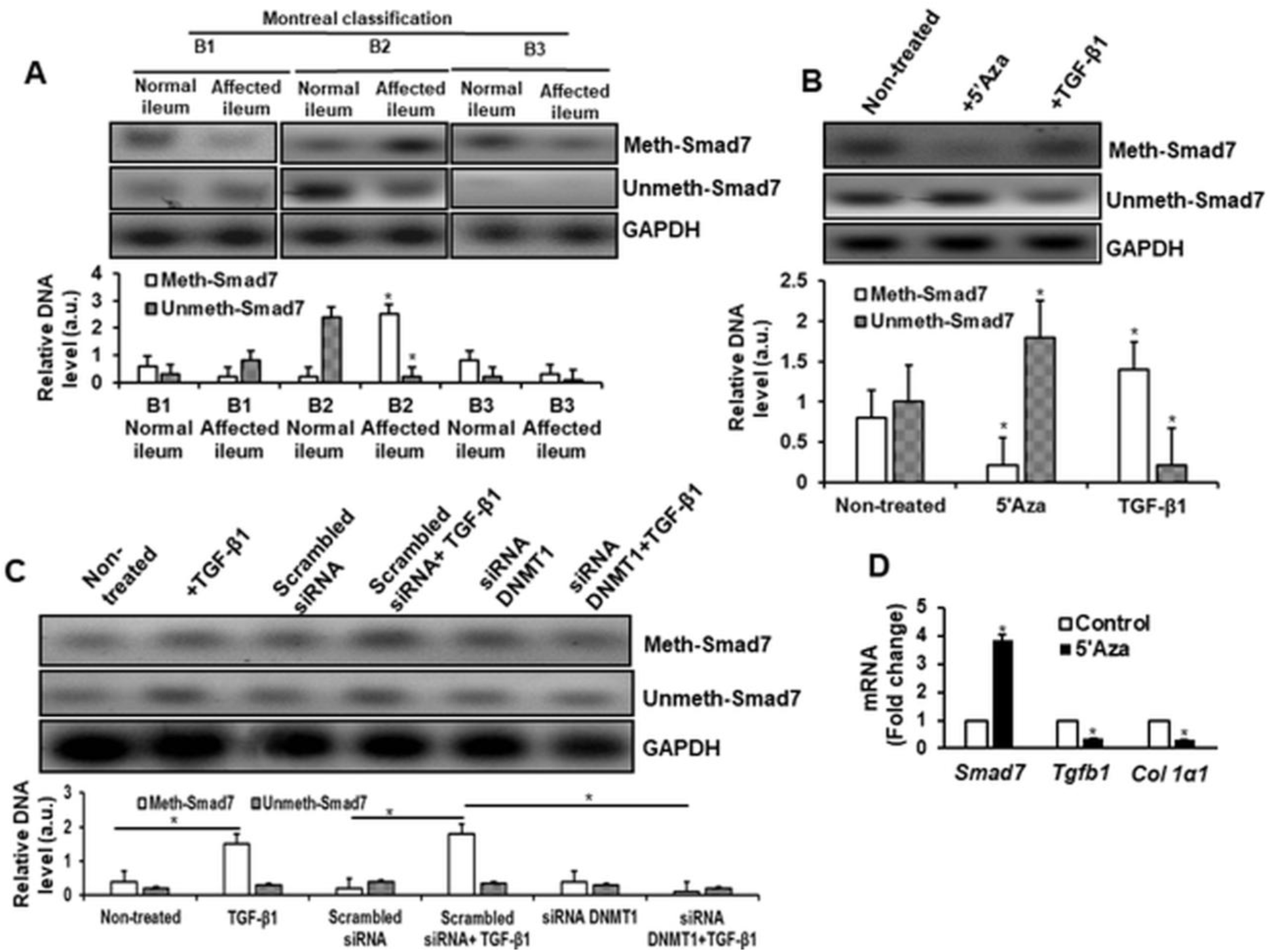
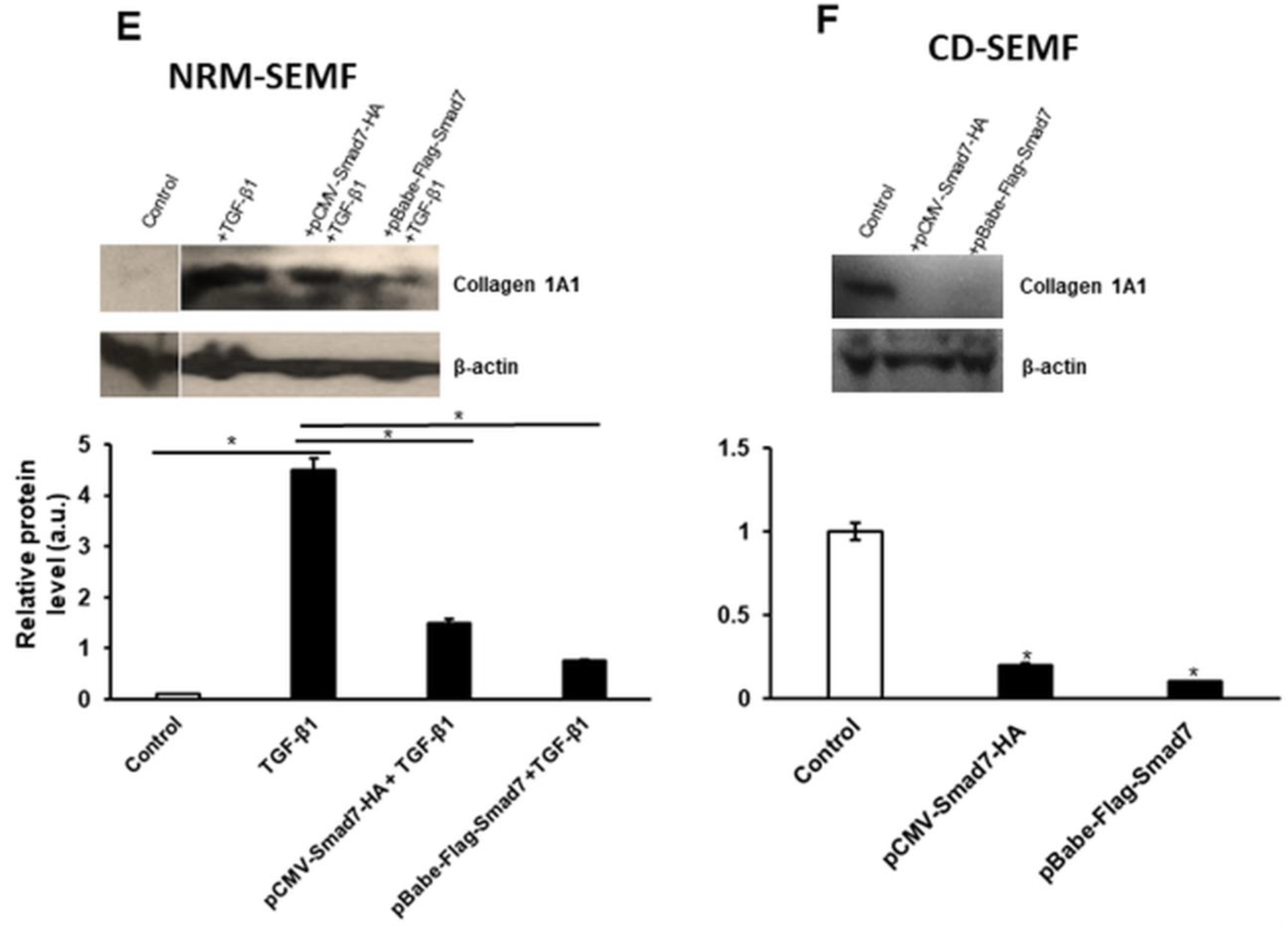
DNMT1-mediated methylation silences Smad7. (A) Representative methylation pcr (upper panel) and densitometric analysis (lower panel) showing increased methylated (□) vs unmethylated (▪) *Smad7* in affected ileum in subjects with B2 fibrostenotic phenotype compared to normal ileum in the same subject. The opposite pattern was seen in patients with B1 inflammatory and B3 penetrating phenotypes. (B) Representative methylation-PCR (upper panel) and densitometric analysis (lower panel) showing TGF-β1 elicits increased methylated (□) vs unmethylated (▪) *Smad7* while 5’-azacytidine decreased methylated (□) vs unmethylated (▪) *Smad7* of SEMF from normal ileum of B2 subjects. (*C*) Representative methylation-PCR (upper panel) and densitometric analysis (lower panel) showing TGF-β1 (10 ng/ml) increased methylated (□) vs unmethylated (▪) *Smad7,* which is mediated by DNMT-1 and was abolished by siRNA mediated knockdown of DNMT-1 but not by scrambled siRNA control. (*D*) Treatment of SEMF with 5’-azacy increased Smad7 expression in SEMF of fibrostenotic ileum in B2 patients. In these cells where Smad7 expression was restored a concomitant decrease in TGF-β1 and Collagen IαI expression was observed. (E) Smad7 overexpression blocks TGF-β1-induced upregulation of collagen 1A1 protein levels in SEMF isolated from normal ileum margin (NRM-SEMF) and (F) endogenous collagen 1A1 protein levels in SEMF isolated from affected ileum in the same patients with B2 fibrostenotic phenotype (CD-SEMF). SEMF were transiently transfected with pCMV-Smad7-HA and pBabe-Flag-Smad7 as well as empty vector overnight then treated with TGF-β1 (10 ng/ml) for additional 16 hours. Cells then were lysed with cell lysis buffer and lysates were subjected to Western blot analysis and quantification analysis of band density was listed below. β-actin was used as a loading control. *P < 0.05 vs normal control, **P < 0.05 vs TGF-β1 treatment. Values represent the mean ± SEM of 4-6 experiments on SEMF from different subjects. Values are expressed in arbitrary units (a.u.).* denotes p < 0.05 vs normal ileum, or non-treated control, or scrambled siRNA, or scrambled siRNA + TGF-β1. Each group data is representative of 6 subjects with the distinct phenotype indicated in the figure.

To determine whether DNMT1 directly regulates SMAD7 methylation, SEMFs isolated from normal ileum of patients with B2 fibrostenotic CD were treated with 5-Azacytidine or transfected with DNMT1-specific siRNA. Both interventions reduced methylated SMAD7 DNA and increased levels of unmethylated SMAD7 DNA (Figure 7B,C). Consistent with these findings, 5-Azacytidine treatment promoted SMAD7 hypomethylation and significantly increased Smad7 mRNA expression (Figure 7D).

To evaluate the functional consequences of restoring Smad7 expression, SEMFs were transfected with two independent Smad7 expression constructs, pCMV-Smad7-HA and pBabe-Flag-Smad7 (Figure 7E,F). Overexpression of Smad7 increased Smad7 protein abundance and was accompanied by reduced collagen Iα1 expression in both SEMFs isolated from normal ileum and affected ileum of the same patients with B2 fibrostenotic CD (Figure 7E,F).

Taken together, these findings demonstrate that DNMT1-mediated promoter hypermethylation contributes to transcriptional silencing of SMAD7 in fibrostenotic Crohn’s disease. Reduced Smad7 expression resulting from promoter hypermethylation represents an additional mechanism that sustains TGF-β1 signaling and promotes intestinal fibrosis.

## DISCUSSION

Our previous work demonstrated that increased TGF-β1 expression and elevated levels of active TGF-β1, resulting in sustained Smad3 activation, are central events in both patients with fibrostenotic CD and mice with chronic TNBS-induced colitis ^3^. These changes promote extracellular matrix production, mesenchymal cell hyperplasia and hypertrophy, and ultimately the development of intestinal fibrostenosis in murine model of colitis ^3^. In the present study, we demonstrate that sustained TGF-β1 signaling and Smad3 activation are maintained through multiple mechanisms that reduce Smad7 abundance and function. First, Smad3 directly regulates pri-hsa-miR-21 transcription, resulting in increased miR-21 expression and degradation of Smad7 mRNA. Second, increased DNMT1 expression promotes hypermethylation and transcriptional silencing of Smad7. In addition, sustained TGF-β1 signaling increases Smurf1/2 activity, resulting in ubiquitination and proteasomal degradation of Smad7. Finally, an altered balance between HDAC1 and p300 activity favors Smad7 deacetylation, thereby enhancing Smurf1/2-dependent ubiquitination and degradation. Together, these mechanisms contribute to persistent suppression of Smad7 and sustained activation of canonical TGF-β signaling in mesenchymal cells of fibrostenotic ileum.

It is noteworthy that many of these regulatory mechanisms also influence Smad7 abundance and function in epithelial cells and T cells; however, they operate differently in those cellular compartments and generally result in increased rather than decreased Smad7 expression^33^.

MicroRNA-21 has been associated with the development of fibrosis in multiple organs^14,55^. Elevated miR-21 expression has previously been reported in mucosal pinch biopsy specimens from patients with active Crohn’s disease^18^. In addition to the Smad3-dependent regulation of pri-miR-21 transcription demonstrated here in mesenchymal cells from fibrostenotic intestine, miR-21 has also been shown to increase intestinal epithelial tight junction permeability in both human intestinal epithelial cells and murine models of colitis ^56,57^. These observations, together with evidence that miR-21 deficiency reduces susceptibility to DSS-induced colitis, suggest that therapeutic targeting of miR-21 may provide benefit in Crohn’s disease.

Expression of TGF-βRII is increased in the intestine of patients with B2 CD (Supplementary Figure 2)^58^. Increased receptor expression may not only facilitate sustained canonical TGF-β signaling but may also occur because of reduced Smad7 expression. Under normal physiological conditions, Smad7 competes with Smad3 for binding to the TGF-β receptor complex and promotes E3 ubiquitin ligase-mediated degradation of TGF-βRI/II complex as part of the negative-feedback mechanism initiated by canonical TGF-β signaling^8,31,32,34^.

Acetylation of Smad7 lysine residues (K64 & K70) is regulated by the balance between p300 acetyltransferase and HDAC1 activity and plays a major role in determining susceptibility to ubiquitination and proteasomal degradation ^59^. In epithelial cells and lamina propria mononuclear cells (LPMC), p300-mediated acetylation stabilizes Smad7 and protects it from degradation, thereby limiting TGF-β1-induced Smad3 activity and supporting regulatory T-cell function ^32,34^. In contrast, the opposite pattern is observed in mesenchymal cells of fibrostenotic intestine, where reduced acetylation targets Smad7 for degradation (Figure 5). The greater increase in HDAC1 relative to p300 observed in SEMFs from affected ileum of patients with B2 fibrostenotic CD shifts the balance toward deacetylation and enhanced Smad7 turnover. Although several acetyltransferases and deacetylases may be involved, the importance of this pathway is supported by the ability of the HDAC inhibitor trichostatin A (TSA) and the p300 inhibitor (C646) to regulate Smad7 protein levels in opposite directions.

The role of Smad7 in CD appears to differ substantially between epithelial and immune cells on one hand and mesenchymal cells on the other. In epithelial cells and LPMC, increased Smad7 expression impairs TGF-β1-dependent suppression of NF-κB activation and disrupts regulatory T-cell function^9,10,33,47^. Knockdown of Smad7 using the antisense oligonucleotide Mongersen (GED-0301) restores Smad3 phosphorylation, alters cytokine production, and improves disease severity in patients with CD and in several murine models of colitis, including TNBS, oxazolone, and SCID models^60,61^. However, the patient populations included in these studies represented mixed CD phenotypes, and fibrosis was not specifically evaluated as a study endpoint ^62^. More recently, a phase III multicenter, randomized, double-blind, placebo-controlled trial reported that GED-0301 was ineffective for the treatment of active CD ^63^. The investigators were unable to determine whether the lack of efficacy reflected inadequate tissue exposure, failure of the intended pharmacodynamic effects, or characteristics of the study population because tissue drug concentrations and histological assessments were not available^63^. Consequently, the role of Mongersen and Smad7-targeted therapies in specific CD phenotypes remains unresolved. Their effects in patients with fibrostenotic disease are unknown. This distinction may be especially important because both human studies and experimental models have demonstrated that intestinal fibrosis can progress despite the absence of ongoing active inflammation ^3,64^.

Downregulation of Smad7 has also been implicated in fibrosis of other organs. Experimental studies have shown that reduced Smad7 expression combined with increased Smurf1/2 activity contributes to renal fibrosis, that Smad7 gene transfer attenuates bleomycin-induced pulmonary fibrosis, and that hepatocyte-specific Smad7 expression protects against liver fibrosis ^65^. These observations support a broader role for Smad7 in fibrogenesis across organ systems and suggest that restoration of Smad7 in a well-phenotyped cohort of patients may represent a therapeutic strategy for a variety of fibrotic disorders.

A better understanding of epigenetic mechanisms regulating key profibrotic factors such as Smad7 may facilitate the development of more precise antifibrotic therapies. Advances in single-cell sequencing, multi-omics technologies, and AI-assisted analyses have uncovered novel molecular and cellular interactions among diverse cell populations involved in fibrosis. Recent single-cell RNA sequencing and spatial transcriptomic studies have substantially advanced our understanding of cellular architecture and unique function of novel differentially expressed genes in multicellular network underlying fibrostenotic CD ^66,67^. Single-cell RNA sequencing of strictured intestinal tissue has revealed marked heterogeneity within mesenchymal populations, including inflammatory fibroblasts and collagen-hi myofibroblasts, and a loss of tissue fibroblasts uniquely associated with fibrotic areas ^66^. These populations of IL-11-producing and high-collagen fibroblasts segregated to distinct areas and expressed cytokines and chemokines in proximity to T or epithelial cells, highlighting inflammatory fibroblasts coordinate local immune response in the gut ^66^. Moreover, Mukherjee et al reported a previously unknown fibroblast heterogeneity and interactions, while CXCL14+ & MMP/WNT5A+ fibroblasts playing an important role in stricutres of Crohn’s disease, with the report of cadherin-11, which is a fibroblast-selective surface molecule, through *in vitro* and *in vivo* studies, validated as a potential therapeutic target ^67^.

Intestinal fibrosis is a complex multicellular process involving interactions among mesenchymal, epithelial, immune, endothelial, and smooth muscle cells. Although the present study focused on SEMF because of their established profibrotic properties, additional cellular populations are likely to contribute to a variety of fibrotic disorders ^68,69^. Future studies will be required to determine whether the mechanisms identified here also regulate Smad7 expression and function in other cell populations involved with fibrosis.

Integrated single-cell and spatial transcriptomic analyses further demonstrate that inflammatory fibroblasts reside within discrete fibrotic cellular neighborhoods together with immune populations, including CCR7^high^ CD4 T cells and IgG-producing plasma cells, suggesting that fibrostenosis arises from coordinated multicellular interactions rather than activation of a single cell type ^66^. These studies also identified spatially organized fibrosis-associated gene programs enriched for extracellular matrix remodeling, TGF-β-responsive pathways, and CD risk loci within fibroblast-rich regions of affected intestine. Particularly notable is the identification of creeping fat-derived CTHRC1-positive fibroblasts as a prominent profibrotic mesenchymal population in CD strictures ^70^. These cells exhibit enhanced extracellular matrix production, mechanosensitive YAP/TAZ signaling, and close association with areas of active fibrosis^70^. Emerging evidence further suggests that a subset of these fibroblasts may originate from creeping fat-associated mesenchymal stromal cells that migrate toward the bowel wall during stricture formation, providing a mechanistic link between mesenteric remodeling and intestinal fibrosis.

Collectively, these findings support a model in which persistent activation of TGF-β/Smad signaling within specialized mesenchymal subsets drives intestinal fibrogenesis. Our findings that Smad7 is suppressed through miR-21-mediated post-transcriptional regulation, DNMT1-dependent promoter hypermethylation, and enhanced ubiquitin-proteasome degradation provide potential molecular mechanisms that may sustain these profibrotic fibroblast phenotypes in fibrostenotic Crohn’s disease. These mechanisms may contribute to the establishment and maintenance of the fibrosis-associated mesenchymal networks recently identified by single-cell and spatial transcriptomic studies.

## Author Contributions

CL: experimental design, acquisition, analysis of data and manuscript writing

JRG: experimental design and manuscript writing, deceased.

KSM: experimental design and manuscript writing, retired.

JB: experimental design and manuscript writing

ER: experimental design and manuscript writing

NW: experimental design and manuscript writing

JFK: Senior Author, experimental design and manuscript writing

